# Oscillatory signal decoding within the ERK cascade

**DOI:** 10.1101/2025.07.24.666680

**Authors:** Ambhighainath Ganesan, Ha Neul Lee, Brian Tenner, Sohum Mehta, Andre Levchenko, Jin Zhang

## Abstract

Extracellular-signal-regulated kinase (ERK) integrates multiple growth factor and hormone stimuli to control essential cellular processes such as proliferation, survival, and migration. In electrically excitable cells, the ERK pathway also interfaces with intracellular Ca^2+^ dynamics to achieve non-canonical, cell-type specific functions, having been implicated in neuronal synaptic plasticity, cardiac hypertrophy, and pancreatic insulin secretion. Yet how the classical Ras/MEK/ERK cascade responds to and decodes dynamic Ca^2+^ signals at its multiple levels to regulate cellular function is poorly understood. Here, we investigated the dynamics of Ca^2+^-induced ERK pathway activation in a pancreatic β-cell line using genetically encoded fluorescent biosensors. By carefully manipulating Ca^2+^ input signals and directly monitoring the activity dynamics of individual ERK pathway components, we reveal that β-cell Ca^2+^ oscillations undergo sequential signal processing along the ERK cascade, mediated by the characteristic response kinetics at each pathway step. We further demonstrate that the ERK cascade and possibly other Ca^2+^-responsive pathways operate within a hybrid network architecture to achieve both hierarchical and parallel processing of β-cell Ca^2+^ oscillations, providing important insights into dynamic signal decoding by this crucial signaling network.

## INTRODUCTION

Signaling pathways are the primary means by which living cells process external information to enable optimal operation of various cellular processes. It is therefore imperative for such pathways to effectively recognize and respond to dynamic changes in the extracellular environment. A classic example of dynamic signaling changes are the oscillatory signals observed across diverse biological contexts (*1*, *2*), from the cell cycle (*3*, *4*) and circadian clock (*5*, *6*) to embryogenesis (*7*) and development (*8*). Biological oscillations can encode information effectively in the temporal/frequency domain (*2*, *9*), yet how signaling molecules respond to these dynamic signals and decode temporal information is not fully understood. Oscillations in intracellular calcium ion (Ca^2+^) concentrations play a particularly key role in driving various processes through temporal encoding of signaling information (*10–13*). Accumulating evidence suggests that target molecules can exhibit different Ca^2+^ response behaviors, with the distinct activation characteristics of individual Ca^2+^ signaling effectors having been shown to yield specific response dynamics, ranging from sustained to fully oscillatory responses, at different Ca^2+^ oscillation frequencies (*14–20*).

Extracellular response kinase (ERK) is a key mediator of various signaling pathways that control essential cellular processes, including proliferation, survival, and migration (*21*, *22*). In electrically excitable cells, ERK also interfaces with Ca^2+^ signaling to achieve myriad cell-type-specific functions. Notably, transcriptional regulation via Ca^2+^-driven ERK signaling plays crucial roles in neuronal synaptic plasticity (*23*, *24*), cardiac hypertrophy (*25–27*), and pancreatic insulin production (*28*). Yet how the classical Ras/MEK/ERK cascade responds to and decodes dynamic Ca^2+^ signals to regulate cellular function is poorly understood. To investigate this question, we turned to pancreatic β-cells as a model of Ca^2+^-dependent ERK signaling.

In β-cells, glucose stimulation triggers robust, cytosolic Ca^2+^ oscillations, which are directly responsible for controlling exocytosis of insulin granules to drive pulsatile insulin secretion by the pancreas (*29*), along with other physiological functions such as gene expression (*30*, *31*) and metabolic homeostasis (*32*, *33*). Glucose, hormones, and membrane-depolarizing agents have all been shown to induce Ca^2+^-dependent ERK activity in β-cells (*34–36*) to regulate insulin gene transcription (*28*), secretory demand (*37*), and overall β-cell health (*38*). The temporal dynamics of ERK activity, especially oscillations (*39*), are thought to influence cell health and are mediated by complex feedback loops in different cells (*40–42*), and previous studies have suggested that ERK maximally responds to certain Ca^2+^ oscillation frequencies (*43*, *44*). Nevertheless, how Ca^2+^ signals are transduced to activate ERK and how oscillatory Ca^2+^ signals are processed along the ERK signaling cascade in pancreatic β-cells remain unclear.

Here, we used genetically encoded fluorescent biosensors to directly visualize the activity dynamics of different components along the ERK signaling cascade in response to oscillatory Ca^2+^ inputs in MIN6 pancreatic β-cells (*45*). We find that kinetic differences at different steps along the ERK cascade play a crucial role in decoding the input Ca^2+^ signal. Sequential processing of the signal by different components of the ERK cascade enables the cell to selectively achieve robust ERK activation while still retaining crucial dynamic information higher up the pathway for use in different contexts.

## RESULTS AND DISCUSSION

To investigate the dynamic control of ERK signaling by Ca^2+^ oscillations, we first used a genetically encoded FRET-based ERK kinase activity reporter, EKAREV (*46*), to visualize ERK activity in real time in live MIN6 β-cells. EKAREV contains an ERK substrate and docking sequence tethered to a WW phosphoamino acid-binding domain via a flexible EV linker. Phosphorylation of the sensor by active ERK results in greater FRET between the flanking fluorescent proteins (FPs) YPet and ECFP, measured as an increase in the yellow-to-cyan (Y/C) emission ratio. To simultaneously capture Ca^2+^ dynamics, we also transfected MIN6 cells with the red-fluorescent Ca^2+^ indicator RCaMP1d (*47*). Glucose has previously been shown to induce Ca^2+^-dependent ERK activation in pancreatic β-cells (*48–50*). Consistent with these observations, we found that treating MIN6 cells with tetraethylammonium chloride (TEA), a potassium channel blocker that mimics the effects of glucose, elicited a robust, Ca^2+^-dependent increase in the EKAREV emission ratio (maximum ratio change, ΔR/R = 42.1 ± 1.3 [mean ± s.e.m.], n = 53 cells; *P* = 3.54 × 10^−36^ vs. −CaCl_2_) (Fig. 1A and Fig. S1). In contrast to the clear Ca^2+^ oscillations reported by RCaMP1d, the EKAREV response showed only minimal fluctuations (Fig. 1A and Fig. S1), revealing the remarkable transformation of an oscillatory Ca^2+^ signal to sustained ERK activity.

**Figure 1.**
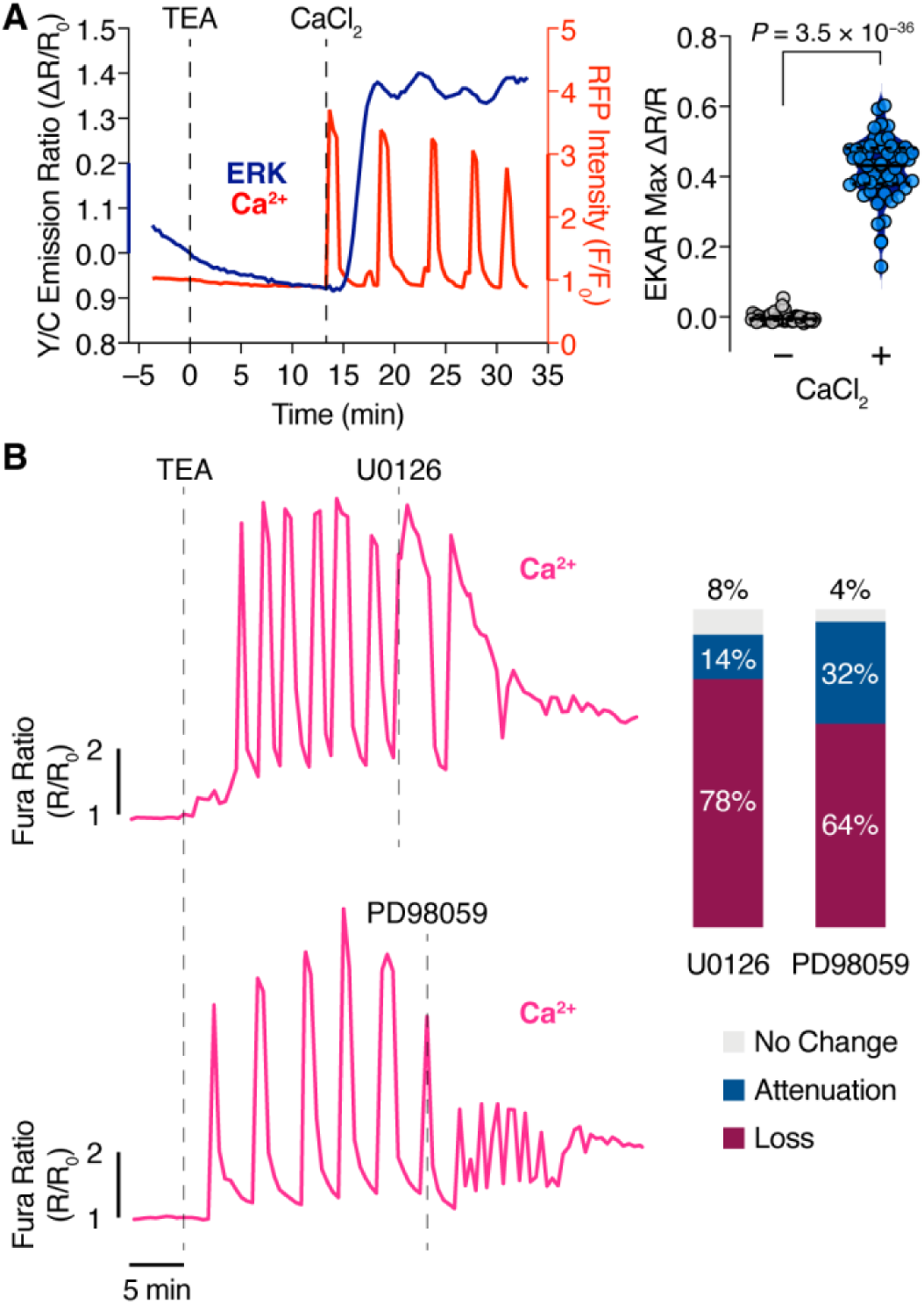
ERK responds to and promotes Ca^2+^ oscillations in MIN6 cells. (A) Left: Representative single-cell timecourse of EKAREV emission ratio (ERK, dark blue curve) and RCaMP1d fluorescence intensity (Ca^2+^, red curve) in MIN6 cells sequentially treated with 20 mM tetraethylammonium chloride (TEA) and 5 mM calcium chloride (CaCl_2_). Curves representative of n = 53 cells from 3 independent experiments. Additional representative traces shown in Fig. S1. Right: Quantification of maximum TEA-stimulated EKAREV emission ratio change (ΔR/R) in MIN6 cells before (−) and after (+) CaCl_2_ addition. n = 53 cells from 3 independent experiments. Data analyzed using paired, two-tailed Student’s t-test. Solid and dashed lines in violin plot show median and quartiles, respectively. (B) ERK pathway inhibition disrupts Ca^2+^ oscillations. Left: Representative single-cell timecourses of Fura-2-stained MIN6 cells sequentially treated with TEA and either 20 μM U0126 (upper) or 20 μM PD98059 (lower) to inhibit MEK. Curves representative of n = 92 (U0126) and 69 (PD98059) cells from 4 experiments each. Additional representative traces are shown in Figs. S2 and S3. Right: Quantification of disrupted TEA-induced Ca^2+^ oscillations following MEK inhibitor treatment in MIN6 cells. Loss: cessation of oscillatory Ca^2+^ dynamics following inhibitor treatment, with Ca^2+^ either remaining steadily elevated or dropping back to baseline levels. Attenuation: continuation of Ca^2+^ oscillations, but with greatly reduced amplitude, after inhibitor treatment.

At the same time, we found that abolishing ERK activation can terminate the oscillatory Ca^2+^ input signal. In MIN6 cells stained with the Ca^2+^ indicator Fura-2 (*51*) and stimulated with TEA, treatment with the MEK inhibitors U0126 (*52*) or PD98059 (*53*) rapidly disrupted Ca^2+^ oscillations in the overwhelming majority of cells (U0126: 85/92 cells; PD98059: 66/69 cells) (Fig. 1B and Fig. S2-3). Most cells exhibited a complete loss of oscillatory behavior after U0126 (79/92 cells) or PD98059 (44/69 cells) treatment, though a subset of cells continued to persist with weaker oscillations under both conditions.

ERK is typically activated by a signaling cascade initiated by the small GTPase Ras (*21*). We therefore investigated Ras activation in MIN6 cells using the FRET-based sensor RaichuEV-Ras (*46*), in which full-length HRas and a Ras-binding domain are joined by an EV linker to report Ras activation via changes in FRET between YPet and the cyan-emitting FP Turquoise-GL. As with EKAREV, TEA stimulation induced a strong, Ca^2+^-dependent emission ratio response from RaichuEV-Ras (ΔR/R = 73.6 ± 4.14%, n = 43 cells; *P* = 2.58 × 10^−20^ vs. −CaCl_2_) (Fig. 2A). MIN6 cells transfected with dominant-negative HRasN17 (*54*, *55*) to block Ras activation also exhibited weaker (ΔR/R = 24.8 ± 1.41%, n = 59 cells) and slower (t_1/2_ = 4.46 ± 0.198 min, n = 59 cells) TEA-induced EKAREV responses than control cells (ΔR/R = 41.8 ± 1.93% n = 58 cells; *P* = 9.28 × 10^−11^; t_1/2_ = 1.48 ± 0.045 min, n = 58 cells; *P* = 2.46 × 10^−34^) (Fig. 2B), confirming that Ras activation is required for Ca^2+^-driven ERK activity in MIN6 β-cells. Ras itself is canonically activated at the plasma membrane by various guanine exchange factors (GEFs) (*56*), including a pair of Ca^2+^-dependent GEFs, RasGRF1/2 (*57*, *58*). We found that siRNA-mediated knockdown of RasGRF2 led to decreased ERK phosphorylation levels in MIN6 cells (Fig. 2C). While multiple signaling molecules have been suggested to activate one or more components of the ERK cascade in β-cells (*34*, *36*, *48*, *59*), our results suggest a role for RasGRF2 in linking Ca^2+^ oscillations to ERK activation in MIN6 β-cells.

**Figure 2.**
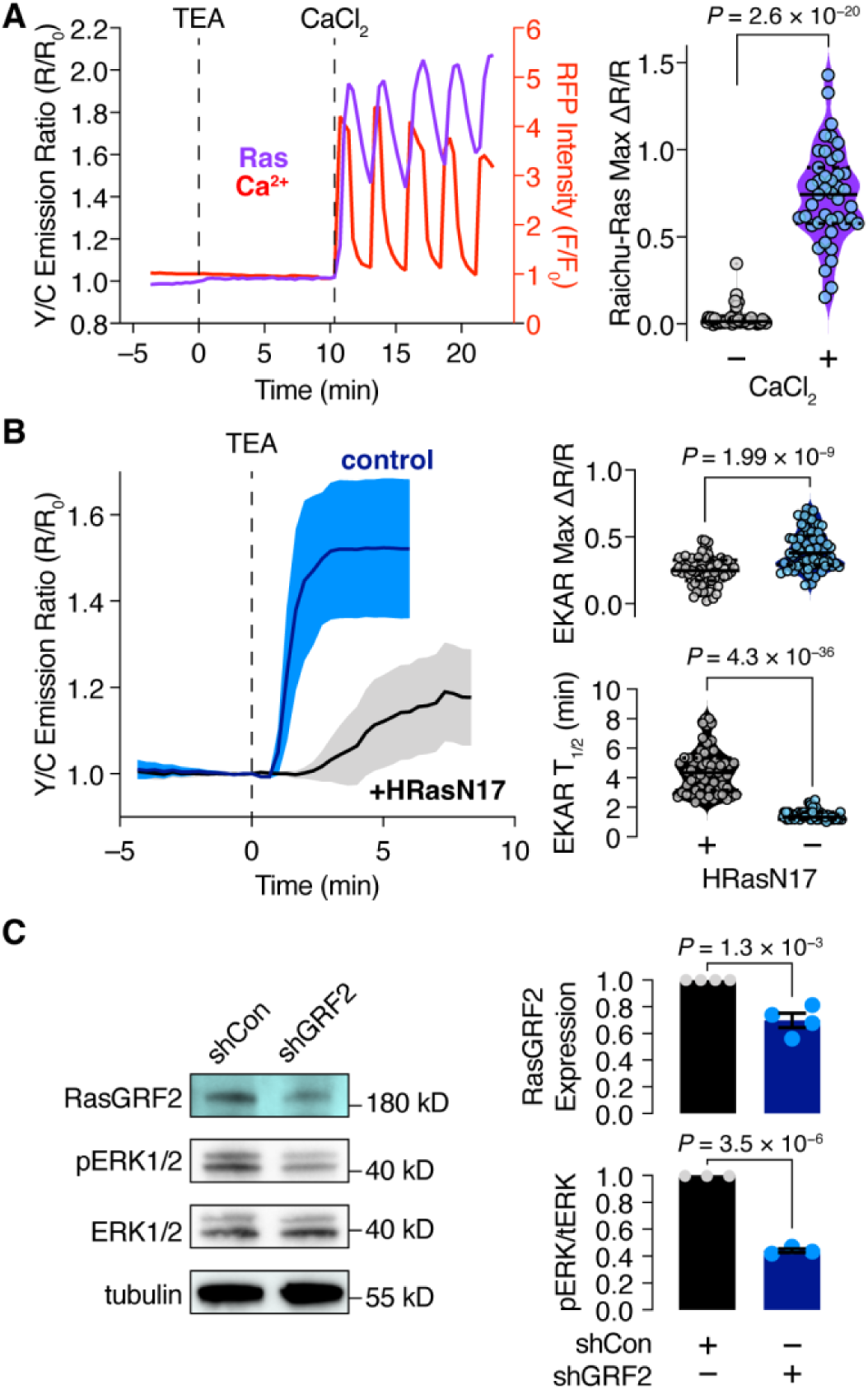
Ca^2+^-dependent ERK activation through Ras and RasGRF2 in MIN6 cells. (A) Left: Representative single-cell timecourse of RaichuEV-Ras emission ratio (Ras, blue curve) and RCaMP1d fluorescence intensity (Ca^2+^, red curve) in MIN6 cells sequentially treated with TEA and CaCl_2_. Curves representative of n = 43 cells from 3 independent experiments. Additional representative traces shown in Fig. S4. Right: Quantification of maximum TEA-stimulated RaichuEV-Ras emission ratio change (ΔR/R) in MIN6 cells before (−) and after (+) CaCl_2_ addition. n = 43 cells from 3 independent experiments. Data analyzed using paired, two-tailed Student’s t-test. (B) Right: Representative average timecourse of EKAREV emission ratio in TEA-stimulated MIN6 cells co-expressing mCherry-NLS (control, dark blue curve; n = 18 cells) or dominant-negative HRasN17 (black curve). n = 18 (control) and 12 (HRasN17) cells, representative of 5 and 7 independent experiments, respectively. Left: Quantification of maximum emission ratio change (ΔR/R; upper) or time to half-maximal response (T1/2; lower) in TEA-stimulated MIN6 cells expressing EKAREV plus either HRasN17 (+) or mCherry-NLS (−). n = 59 (+) and 65 (−) cells from 5 and 7 independent experiments, respectively. Data analyzed using Mann-Whitney U-test. Solid and dashed lines in violin plots show median and quartiles, respectively. (C) Representative western blot (left) and quantification of RasGRF2 expression (upper right; n = 4 independent experiments) and ratio of phosphorylated ERK to total ERK expression (pERK/tERK; lower right; n = 3 independent experiments) in MIN6 cells expressing either control shRNA (shCon) or shRNA targeting RasGRF2 (shGRF2). Data analyzed using unpaired, two-tailed Student’s t-test. Bars show mean ± s.e.m (normalized to shCon).

We next set out to investigate how MIN6 β-cells convert the oscillatory Ca^2+^ input signal into sustained ERK activity. Highly dynamic control of cellular processes can be achieved by minimizing kinetic differences between the input signal and output response, while conversely pairing a dynamic input with a slow output could allow integration of input fluctuations (Fig. 3A). Thus, the deactivation kinetics (i.e., the time to reach baseline levels after stimulation) of a given signaling component should dictate its ability to dynamically react to input changes such as Ca^2+^ oscillations. Borrowing a well-established concept in control theory (*60*), we quantified the impulse responses of each pathway component (i.e., RasGRF2, Ras, ERK) following a single input Ca^2+^ “pulse” induced by transient depolarization using potassium chloride (KCl) (Fig. 3B). Strikingly, we observed that the deactivation kinetics became progressively slower at each step of the ERK pathway (Fig. 1C-F). Using mCherry-tagged RasGRF2, which translocates to and from the plasma membrane in response to RasGRF2 activation, we measured RasGRF2 deactivation kinetics (t_1/2(decay)_ = 0.687 ± 0.021 min [mean ± s.e.m.], n = 16 cells) that closely tracked with the input Ca^2+^ signal (t_1/2(decay)_ = 0.745 ± 0.021 min, n = 109 cells; *P* = 0.256, Mann-Whitney U-test) (Fig. 1C). In contrast, RaichuEV-Ras reported Ras deactivation kinetics on the order of ∼7 min (t_1/2(decay)_ = 7.35 ± 0.562 min, n = 32 cells; *P* = 8.87 × 10^-13^ vs. RasGRF2). Finally, EKAREV imaging revealed that downstream ERK activity took ∼11 min on average (t_1/2(decay)_ = 11.2 ± 0.426 min, n = 70 cells; *P* = 6.22 × 10^-8^ vs. Ras; Mann-Whitney U-test) to decay back to half-maximal levels in response to a Ca^2+^ pulse (Fig. 1D-F). Furthermore, deactivation of the ERK response could be accelerated using the MEK inhibitor U0126 in conjunction with washout of KCl (Fig. S2), suggesting that the slow kinetics are not an artifact of the EKAREV sensor.

**Figure 3.**
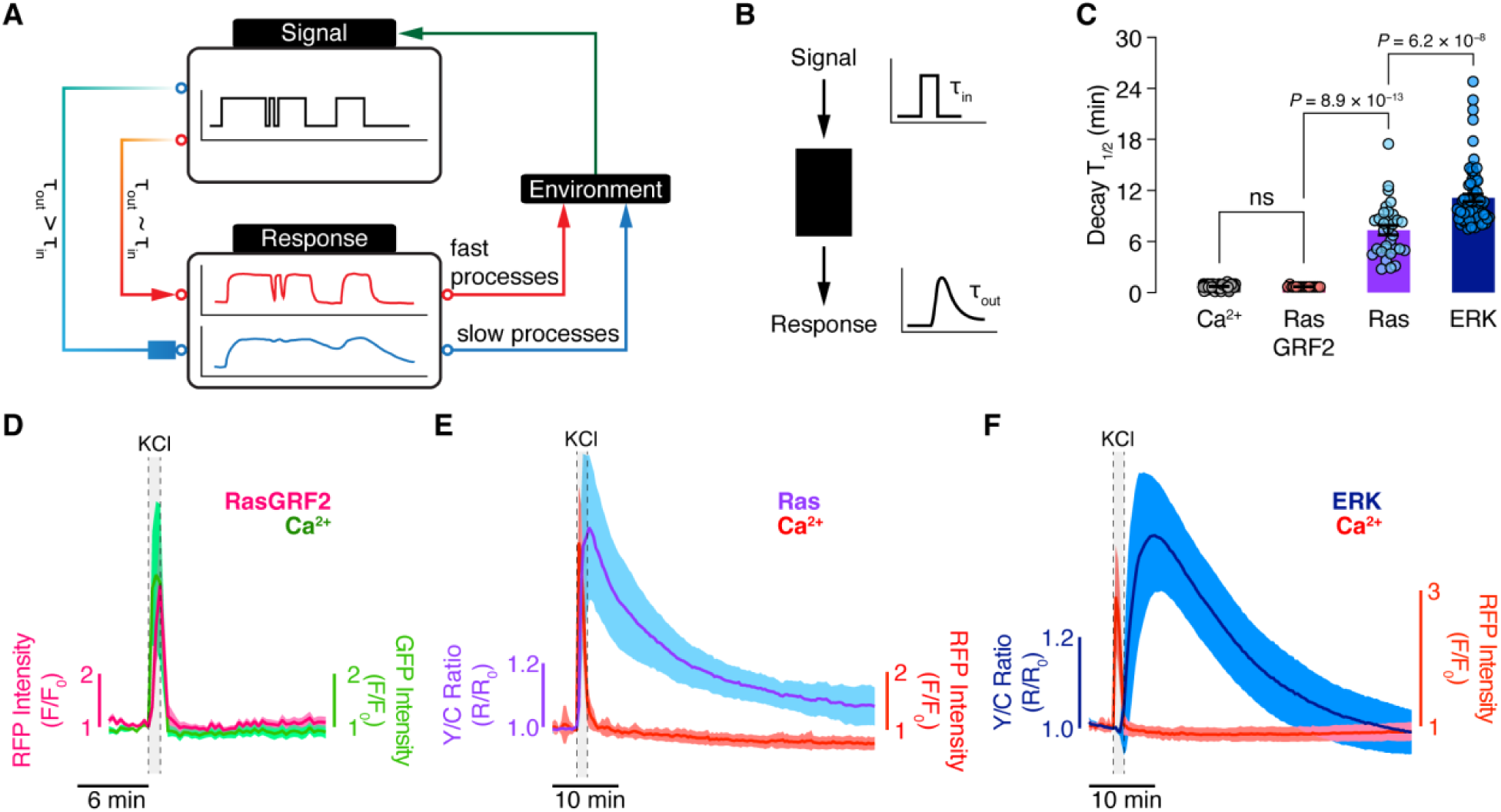
Impulse response dynamics of ERK pathway components in MIN5 cells. (A) Diagram illustrating how the relationship between input (τ_in_) and output (τ_out_) kinetics influences the processing of extracellular signals into dynamic or sustained intracellular responses to control fast or slow processes that ultimately drive adaptation to environmental changes. (B) Schematic of an impulse-response assay, in which a single, pulsed input is applied to a given system to characterize how the kinetics of a measured response output (τ_out_) vary with respect to the kinetics of the applied input signal (τ_in_). (C) Summary quantification of the time to half decay (Decay T_1/2_) of the response from RCaMP1d (Ca^2+^;), RasGRF2-mCherry (RasGRF2; n = 16 cells), RaichuEV-Ras (Ras; n = 32 cells) and EKAREV (ERK; n = 70 cells) in MIN6 cells simulated with 30 mM potassium chloride (KCl) for 1 min. n = 109 (Ca^2+^), 16 (GRF2), 32 (Ras), and 70 (ERK) cells from 7, 4, 3, and 4 independent experiments, respectively. ns, not significant. Data analyzed using Mann-Whitney U-test. Bars show mean ± s.e.m. (D-F) Representative average timecourses in MIN6 cells co-expressing (D) RasGRF2-mCherry (magenta curve; n = 7 cells) plus pmGCaMP3 (Ca^2+^, green curve) or either (E) RaichuEV-Ras (Ras, blue curve; n = 15 cells) or (F) EKAREV (ERK, dark blue curve; n = 17 cells) plus RCaMP1d (Ca^2+^, red curves) and stimulated with 30 mM KCl for 1 min. Solid lines indicate the mean; shaded areas, s.d. Representative of 4 (D), 3 (E), and 4 (F) independent experiments. See also Fig. S5.

We further observed that RasGRF2, whose deactivation kinetics closely mirrored those of the input Ca^2+^ signal (Fig. 3C, D), rapidly oscillates in tandem with Ca^2+^ in TEA-stimulated MIN6 β-cells (Fig. 4A and S6A). However, Ras and ERK, which exhibited progressively slower deactivation kinetics (∼7 and ∼11 min, respectively) compared with RasGRF2, correspondingly displayed increasingly sustained activity patterns in response to the same TEA-induced oscillatory Ca^2+^ signal (Fig. 4A and S6B, C). These observations suggest that the increasing mismatch in deactivation kinetics at each step of the ERK cascade enables the gradual transformation of an oscillatory Ca^2+^ signal into robust, sustained ERK activity. Notably, the downstream ERK targets c-fos and c-jun, the so-called Immediate Early Gene (IEG) products, can exhibit even slower deactivation kinetics, on the order of a few hours (*52*, *61*, *62*), suggesting that the Ca^2+^-dependent ERK signal in MIN6 cells likely undergoes additional processing before functional targets are activated.

**Figure 4.**
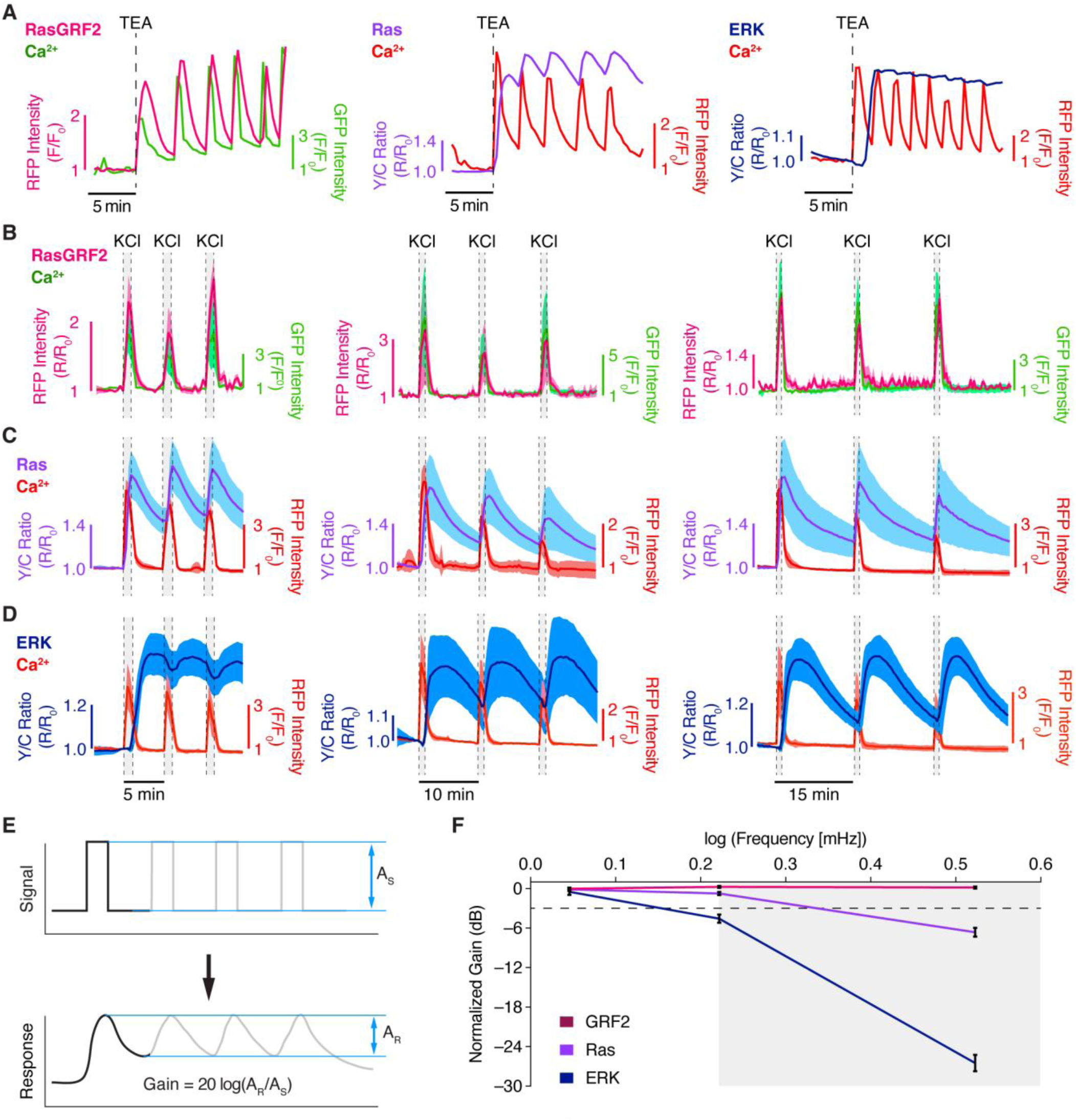
Progressive temporal decoding of Ca^2+^ oscillations down the ERK cascade. (A) Representative single-cell timecourses of TEA-stimulated biosensor responses in MIN6 cells co-expressing (left) RasGRF2-mCherry (magenta curve) plus pmGCaMP3 (Ca^2+^, green curve) or either (middle) RaichuEV-Ras (Ras, blue curve) or (right) EKAREV (ERK, dark blue curve) plus RCaMP1d (Ca^2+^, red curves). Traces are representative of n = 24 (RasGRF2), 20 (Ras), and 14 (ERK) cells from 4, 4, and 3 independent experiments, respectively. Additional representative traces shown in Fig. S6. (B-D) Representative average timecourses of biosensor responses from MIN6 cells co-expressing (B) RasGRF2-mCherry plus pmGCaMP3 (left-to-right, n = 5, 9, and 7 cells) or either (C) RaichuRas-EV (left-to-right, n = 10, 11, and 7 cells) or (D) EKAREV (left-to-right, n = 12, 9, and 15 cells) plus RCaMP1d and stimulated with 30 mM KCl for 1 min at 5- (left), 10- (middle), or 15 (right)-min intervals. Solid lines indicate the mean; shaded areas, s.d. Data are representative of (B) n = 15, 19, and 17 cells from 4, 3, and 4 independent experiments, (C) n = 48, 42, and 35 cells from 6, 4, and 5 independent experiments, and (D) n = 49, 39, and 48 cells from 4 independent experiments each. (E) We assessed the frequency response of each pathway component by measuring the amplitude, defined as the peak-to-trough difference for a single pulse, of the input signal (e.g., Ca^2+^) and output response (e.g., RasGRF2, Ras, or ERK) and calculating the gain at each KCl pulse frequency. (F) Plot showing the normalized gain with respect to the log of the KCl pulse frequency for RasGRF2 (GRF2, magenta), Ras (blue), and ERK (dark blue). Values calculated from data in B-D. Lines indicate the mean; error bars, s.e.m.

A key feature of biological oscillations is their ability to encode information in the temporal/frequency domain. We therefore investigated whether and how the ERK cascade decodes temporal information by monitoring RasGRF2, Ras, and ERK activity dynamics in MIN6 cells stimulated with precise trains of Ca^2+^ pulses via transient KCl application at three different input frequencies – low (15-min interval), moderate (10-min interval) and high (5-min interval) (Fig. 4B-D). MIN6 cells expressing RasGRF2-mCherry exhibited fully oscillatory changes in plasma membrane fluorescence intensity at all three Ca^2+^ input frequencies (Fig. 4B), indicating fully dynamic RasGRF2 activity. At the Ras level, RaichuEV-Ras responses exhibited partial oscillations in response to the high-frequency input and showed more complete oscillatory responses at the moderate and low Ca^2+^ pulse frequencies (Fig. 4C). Finally, EKAREV displayed oscillatory response behavior only at the lowest input frequency (Fig. 4D). Thus, each molecule in the pathway decodes input frequency information distinctly, according to its response time characteristics.

An important concern related to pathway or network signal processing is the inevitable accumulation of noise during sequential signal transduction steps. Since both signal and noise can be present at all different possible input frequencies, it is not clear how noise filtering might occur without affecting the signal content. Significantly, the progressive increase in deactivation kinetics over sequential pathway steps suggests that the capacity to filter out high frequency signals will differ at different levels along the cascade. To quantitatively test this hypothesis, we measured the “gain” (i.e., the capacity to transmit oscillatory information), defined as the ratio of the output and input signal amplitudes, at each step of the cascade (Fig. 4E). When the output module faithfully reproduces an oscillatory input signal, the system has high gain. Conversely, low gain indicates processing of an oscillatory signal to a sustained output. Indeed, we found that RasGRF2 possesses consistently high gain at all frequencies tested (Fig. 4F), indicating a high “bandwidth” for faithfully transmitting oscillatory information. Downstream signaling components like Ras and ERK, on the other hand, which exhibited slower deactivation kinetics, displayed correspondingly lower gain, resulting in the integration of oscillatory signals at higher frequencies (Fig. 4F).

From our results, the ERK signaling cascade in MIN6 cells appears to act as a “low-pass filter”, similar to homologous pathways in worms (*44*) and NIH3T3 cells (*63*) but in contrast to the yeast ERK homolog Hog-1, which was shown to act as a band-pass filter via negative feedback (*64*, *65*). Yet despite the gradual filtering of high-frequency signals down the cascade, we hypothesized that this pathway may still be able to utilize dynamical information through the activation of other signaling molecules by upstream components (Fig. 5A). For instance, Ca^2+^ can directly activate other signaling pathways in parallel to the RasGRF/Ras/ERK cascade, such as PKA (*66–69*) and PKC (*70*, *71*), while RasGRF2 is an activator of not only Ras but also the small GTPase Rac1 (*72*, *73*). Indeed, the FRET-based PKA sensor AKAR4 (*74*) showed highly dynamic, oscillatory responses in TEA-stimulated MIN6 cells (Fig. 5B and Fig. S7A), consistent with our previous observations (*66*, *67*). PKC activity, visualized using the FRET-based sensor CKAR2 (*75*), showed similarly oscillatory response dynamics following TEA stimulation (Fig. 5C and Fig. S7B). Both PKA and PKC activity thus fully retained oscillatory information, consistent with their proximity to the Ca^2+^ input. Meanwhile, RaichuEV-Rac1 (*46*) exhibited responses that were more variable than those displayed by RaichuEV-Ras, which frequently followed the input without integrating it, suggesting faster signal decoding vs Ras (Fig. 5D and S7C). Such fast oscillatory information may likely be used for other functional processes such as insulin secretion via Rac1-mediated actin remodeling (*76*) that may need to occur on a faster time scale than that of the Ras dynamics. These results support our hypothesis that dynamical information is functionally retained early in the cascade, while being simultaneously processed into a robust, downstream response.

**Figure 5.**
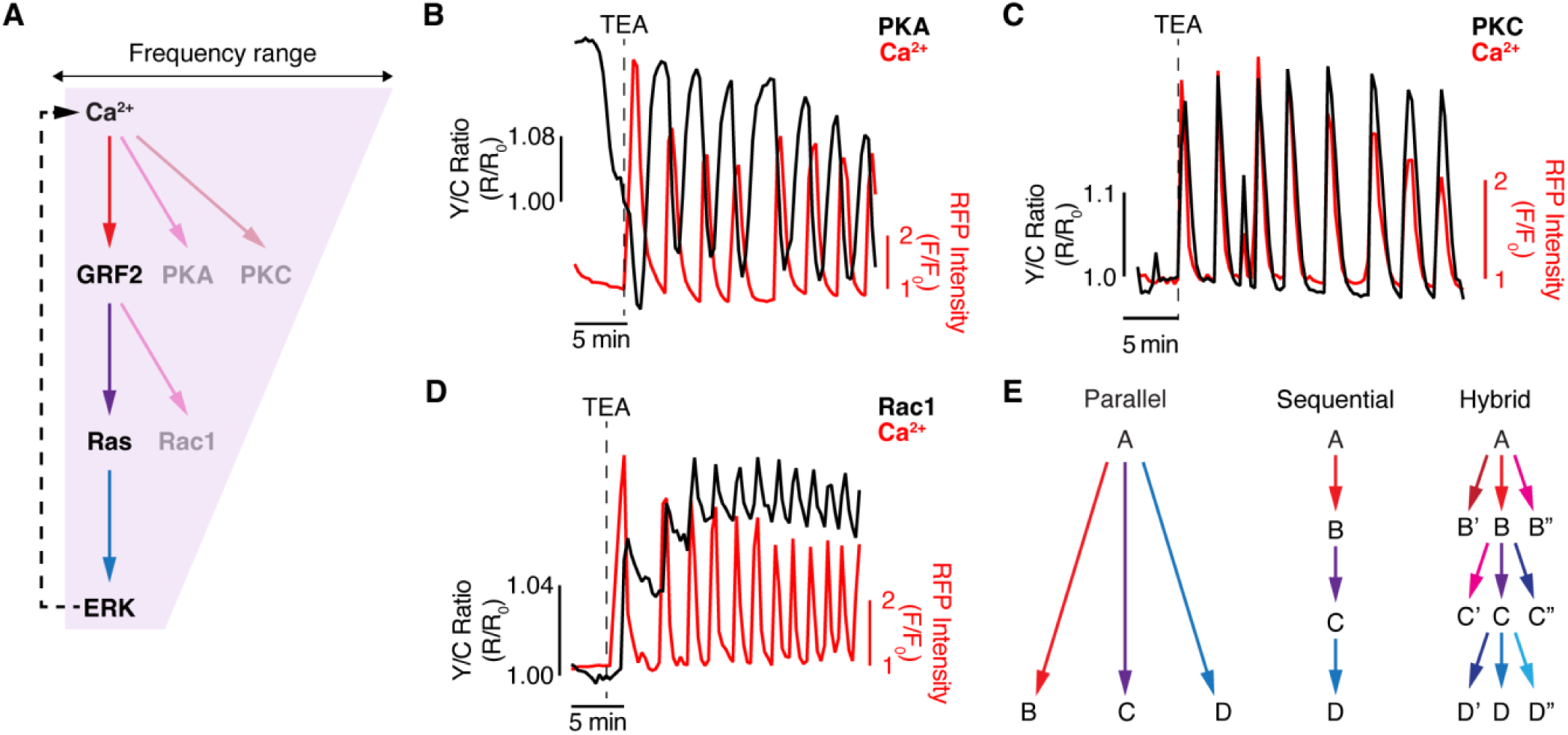
A hybrid network processes oscillatory Ca^2+^ signaling in MIN6 cells. (A) Simplified schematic of the ERK cascade in MIN6 cells. Vertical arrows indicate information flow down the cascade. Branching arrows highlight additional targets that can be activated by upstream pathway components. The dashed arrow indicates feedback regulation of the Ca^2+^ signal by ERK activation. At each step of the cascade, temporal information is filtered out, decreasing the frequency range (shading). (B-D) Representative single-cell timecourses showing TEA-stimulated emission ratio (black curves) and fluorescence intensity (red curves) responses in MIN6 cells co-expressing (B) the PKA sensor AKAR4, (C) the PKC sensor CKAR2, or (D) the Rac1 sensor RaichuEV-Rac1 plus the Ca^2+^ sensor RCaMP1d. Data are representative of (B) n = 21, (C) 26, and (D) 19 cells from 6, 3, and 5 independent experiments, respectively. Additional representative traces are shown in Fig. S7. (E) Our data suggest that the ERK cascade in MIN6 cells combines the parallel signal processing of a “fan-like” signaling network (left) with the sequential processing of a hierarchical cascade (middle) into a hybrid signaling network architecture (right) to decode oscillatory signaling inputs into robust and diverse cellular responses.

Parallel decoding along a “fan-like” network that links discrete downstream pathways (Fig. 5E, left) can be a convenient means for a dynamic input to control multiple cellular functions (e.g., through stimulation of distinct gene subsets at different input frequencies). Alternatively, sequential decoding along a “hierarchical” cascade can impose vital checkpoints in the transfer of information to filter out input noise (Fig. 5E, middle). Our work reveals that the ERK pathway in MIN6 cells comprises a “hybrid” network that combines the hierarchical behavior of a signaling cascade with the egalitarian features of fan-like networks to decode oscillatory Ca^2+^ signals (Fig. 5E, right). This architecture is characterized by fast responses that may have shorter memory at the signaling steps close to the signaling input (e.g., for RasGRF2), thus ostensibly implying the more immediate responses at these levels, not necessarily modulated by the prior history of the signal exposure. However, the feedback from the more downstream, slower response and greater memory steps can establish such prior exposure context. Indeed, our results demonstrate existence of such feedback, enabling the flow of time-integrated information from ERK to the upstream Ca^2+^ control step and ensuring that maximal ERK activation serves a crucial checkpoint for reliable transmission of oscillatory Ca^2+^ information. This type of signaling network architecture enables both the faithful transmission of relevant input information and filtering of high-frequency noise at the level of functionally active downstream targets while ensuring that components higher up the cascade retain the ability to transmit crucial, high-frequency oscillatory information across different pathway contexts. Other pathways may exhibit similar input decoding logic, as recently suggested for the network downstream of VEGF (*77*). Indeed, our previous work revealed that activation of the phosphatase calcineurin (CaN) closely tracks oscillatory Ca^2+^ inputs in MIN6 cells, while its downstream target nuclear factor of activated T-cells (NFAT) (*78*) responds on a slower timescale, leading to stepwise integration of the input dynamics (*79*). Fast CaN activation dynamics are likely essential for regulating targets such as ion channels (*80*) but dispensable for the slower transcriptional regulation by NFAT. Overall, it is tempting to consider that gradually decreasing response bandwidth might be a common feature of many signaling networks, enabling a graceful tradeoff between response kinetics and noise accumulation in multi-step signaling networks.

## MATERIALS AND METHODS

### Plasmids

AKAR4 (*74*) and CKAR2 (*75*) were described previously. EKAREV, RaichuEV-Ras, RaichuEV-Rac1 (*46*), and pERedNLS-HRasN17 (*81*) were generously provided by Michiyuki Matsuda (Kyoto University). RCaMP1d (*47*) was provided by Loren Looger (UC San Diego). RasGRF2-mCherry was a gift of Mike Moran (MDS Proteomics). pmGCaMP3 was generated by PCR amplification of GCaMP3 (*82*) (gift of Loren Looger, UC San Diego), followed by ligation into a *Bam*HI/*Eco*RI-digested pcDNA3 (Invitrogen) backbone containing the C-terminal KKKKKSKTKCVIM sequence from KRas. All plasmids were verified by Sanger sequencing (Azenta).

### Cell culture and transfection

MIN6 β-cells (*45*) (gift of Jun-Ichi Miyazaki, Osaka University) were cultured in DMEM (Gibco) containing 4.5 g/L glucose and supplemented with 10% (vol/vol) FBS, 1% (vol/vol) penicillin-streptomycin, and 50 μM β-mercaptoethanol and maintained at 37 °C in a humidified, 5% CO_2_ atmosphere. Prior to transfection, cells were plated onto sterile, 35-mm glass-bottom dishes and grown for 48 h to reach approximately 50% confluence. Cells were then transfected using Lipofectamine 2000 (Invitrogen) and grown an additional 48 h before imaging. For Fura-2 imaging, untransfected MIN6 cells (∼70-80% confluence) were loaded with 1 μM Fura-2-acetoxymethylester (Fura-2-AM, Molecular Probes) for 10-20 min at 37 °C.

### Generation of stable RasGRF2 knockdown MIN6 β-cells

shRNA sequences targeting mouse RasGRF2 were designed using the BLOCK-iT™ RNAi Designer (Thermo Fisher Scientific; https://rnaidesigner.thermofisher.com). Double-stranded oligonucleotides were prepared by annealing complementary pairs of synthesized DNA oligonucleotides. The annealed oligonucleotides were cloned into *Age*I/*Eco*RI-digested pLKO.1 vector. For lentivirus production, HEK293T cells were transfected with pLKO.1 constructs along with packaging plasmids (psPAX2:pMD2.G:pLKO.1 at a 9:1:10 ratio) using Lipofectamine™ 2000 (Invitrogen) according to manufacturer’s instructions. Viral supernatants were collected 48 h post-transfection and used to infect MIN6 β-cells in the presence of polybrene (10 μg/ml).

After 48 h, stably transduced cells were selected with puromycin (1 μg/ml) for 3 days. Complete cell death in untransduced control cells confirmed successful selection. The sequences of shRNA oligonucleotides were as follows: control shRNA (forward: 5’-CCGGCAACAAGATGAAGAGCACCAACTCGAGTTGGTGCTCTTCATCTTGTTGTTTTT-3’, reverse: 5’-AATTAAAAACAACAAGATGAAGAGCACCAACTCGAGTTGGTGCTCTTCATCTTGTTG −3’) and RasGRF2 shRNA (forward: 5’-CCGGGCGCGATAGCGCTAATAATTTCTCGAGAAATTATTAGCGCTATCGCGCTTTTT-3’, reverse: 5’-AATTAAAAAGCGCGATAGCGCTAATAATTTCTCGAGAAATTATTAGCGCTATCGCGC-3’).

### Western blotting

Protein lysates were prepared from MIN6 β-cells by washing twice with PBS followed by lysis in ice-cold RIPA buffer (140 mM NaCl, 10 mM Tris-Cl pH 8.0, 1% Triton X-100, 0.1% SDS, 0.1% sodium deoxycholate, 1 mM EDTA, 0.5 mM EGTA) supplemented with protease inhibitor cocktail (Roche Applied Science). After centrifugation (15 min, 12,000 rpm, 4°C), protein concentrations were determined using a Pierce™ BCA assay kit (Thermo Fisher Scientific). Equal amounts of protein were resolved on 4-20% Mini-PROTEAN® TGX™ gels (Bio-Rad) and transferred to PVDF membranes (Millipore). The membranes were blocked with 5% skim milk in TBST (0.1% Tween-20) for 30 min at room temperature, then probed overnight at 4°C with primary antibodies against RasGRF2 (Abcam, ab226973; 1:1000), phospho-ERK1/2 (Cell Signaling Technology #9106S; 1:1000), total ERK1/2 (Cell Signaling Technology #4695P; 1:1000), or β-tubulin (Cell Signaling Technology #2146S; 1:5000). After washing in TBST, membranes were incubated with appropriate HRP-conjugated secondary antibodies (1:5000) for 1 h at room temperature. Immunoreactive bands were visualized using SuperSignal™ West Femto (Thermo Scientific) chemiluminescent substrate on a ChemiDoc™ Imaging System (Bio-Rad) and quantified using ImageJ software.

### Time-lapse fluorescence imaging

#### Widefield epifluorescence imaging

MIN6 cells were washed twice with HBSS and subsequently imaged in HBSS in the dark at 37 °C. Tetraethyammonium chloride (TEA; Sigma), calcium chloride (CaCl_2_; Fisher), potassium chloride (KCl; J.T. Baker), U0126 (Sigma), and PD98059 (Cayman Chemical) were added as indicated.

Live-cell imaging of Fura-2-stained cells was performed on a Zeiss Axiovert 200M (Carl Zeiss) equipped with a 40×/1.3 NA oil objective and a MicroMAX BFT512 cooled charge-coupled device camera (Roper Scientific, Trenton, NJ) controlled by METAFLUOR 7.7 software (Molecular Devices). Dual Fura-2 excitation-ratio imaging was performed using 350DF10 and 380DF10 excitation filters, a 450DRLP dichroic mirror, and a 535DF45 emission filter. Filter sets were alternated using a Lambda 10-2 filter-changer (Sutter Instruments). Exposure times were 500 ms for each channel, and images were acquired every 30 s.

Multiplexed live-cell imaging of RCaMP1d plus EKAREV, RaichuEV-Ras, AKAR4, CKAR2, or RaichuEV-Rac1 was performed on a Zeiss AxioObserver Z7 microscope (Carl Zeiss) equipped with a Definite Focus.2 system (Carl Zeiss), a 40×/1.4 NA oil objective, and a Photometrics Prime95B sCMOS camera (Photometrics) and controlled by METAFLUOR 7.7 software (Molecular Devices). Dual cyan/yellow emission ratio imaging was performed using a 420DF20 excitation filter, a 455DRLP dichroic mirror and two emission filters (473DF24 for CFP and 535DF25 for YFP). RFP intensity was imaged using a 572DF35 excitation filter, a 594DRLP dichroic mirror, and a 645DF75 emission filter. Filter sets were alternated by an LEP MAC6000 control module (Ludl Electronic Products Ltd.). Exposure times ranged from 50 to 500 ms, and images were acquired every 20 s.

#### Total internal reflection fluorescence (TIRF) imaging

The membrane translocation of RasGRF2-mCherry was visualized using total internal reflection fluorescence (TIRF) microscopy on an Olympus IX83 microscope equipped with a cellTIRF-4Line system, a 150× NA 1.45 objective and an Andor IXON Ultra 888 electron-multiplying CCD camera. For imaging, cells were maintained in HBSS and stimulated with 20 mM TEA. Images were acquired using a Cell* 561 nm laser in TIRF mode, with the TIRF angle adjusted to optimize the visualization of plasma membrane-localized RasGRF2-mCherry. Time-lapse images were captured every 20 sec.

#### Image analysis

Raw fluorescence images were analyzed using METAFLUOR 7.7 (Molecular Devices) or ImageJ (FIJI) (*83*) software. Regions of interest (ROI) were drawn around biosensor-expressing or Fura-2-stained cells, along with a cell-free region (background), and background-subtracted fluorescence intensities were calculated by subtracting the fluorescence intensity of the background ROI from the emission intensities of the cell ROIs. Yellow-over-cyan emission ratios and Fura-2 excitation ratios were then calculated at each time point. Response time-courses were plotted as the normalized emission or excitation ratio (R) or fluorescence intensity (F) with respect to time zero (e.g., R/R_0_ or F/F_0_), where R and F are the ratio or intensity value at a given time point, and R_0_ and F_0_ are the initial ratio or intensity value immediately preceding drug addition. Maximum ratio (ΔR/R) changes were calculated as (R_max_−R_min_)/R_min_ or (F_max_−F_min_)/F_min_, where R_max_ and R_min_ or F_max_ and F_min_ are the maximum and minimum ratio or intensity value, respectively, recorded after drug addition. Time to half-maximal response (T_1/2_) and time to half-maximal decay (decay T_1/2_) values were determined from single-cell traces by manually identifying the timepoint after T_0_ (for T_1/2_) or T_peak_ (for decay T_1/2_) with a ratio value nearest to the calculated half-maximal ratio (ΔR/2) or intensity (ΔF/2) change. Linear interpolation was performed when the half-maximal response value fell between two timepoints. For KCl-pulse-train experiments, gain (dB) was calculated using the formula

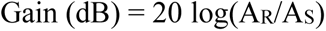

where A_S_ represents the amplitude of the input Ca^2+^ signal, measured as the difference between the maximum normalized GCaMP or RCaMP1d fluorescence intensity (F_max_) reached after a given KCl pulse and the minimum intensity (F_min_) reached immediately before application of the next KCl pulse, and A_R_ represents the amplitude of the output response, measured as above based on RasGRF2 fluorescence intensity (F_max_-F_min_) or RaichuEV-Ras or EKAREV emission ratio (R_max_-R_min_). TEA-induced Ca^2+^ responses in MIN6 cells treated with MEK inhibitors were manually classified as “No Change”, “Attenuation” or “Loss”. Investigators were not blinded to the treatment conditions. All graphs were plotted in GraphPad Prism 10 (GraphPad Software).

### Statistics and reproducibility

All experiments were independently repeated as indicated in the figure legends. All replication attempts were successful. Statistical analyses were performed using GraphPad Prism 10 (GraphPad Software). Pairwise statistical comparisons were performed using paired or unpaired Student’s t-test (Gaussian data) or the Mann-Whitney U test (non-Gaussian data) as indicated. Statistical significance was set at P < 0.05. Unless otherwise noted, averaged live-cell imaging time-courses depict mean ± s.d., violin plots depict the median and quartiles, and bar graphs show mean ± s.e.m.

### Data availability

Plasmids generated in this study will be made available through Addgene (www.addgene.org/Jin_Zhang). Source data are provided with the manuscript.

### Code availability

No custom code was used in this study.

## ACKNOWLEDGEMENTS

The authors thank Michiyuki Matsuda (Kyoto University) and Loren Looger (UC San Diego) for plasmids and Jun-Ichi Miyazaki (Osaka University) for cell lines. We are also grateful to Wei Lin for assistance with TIRF imaging. This work was supported by NIH grants R35 CA197622 and R01 DK073368 (to J.Z.) and R01 GM123011 (to A.L.).

## AUTHOR CONTRIBUTIONS

AG, AL, and JZ conceived of the project. AG, HNL, BT, and SM performed experiments and analyzed the data. JZ, SM, and AL supervised the project. AG, SM, AL, and JZ wrote the manuscript.

