## Supplementary Figures for "Oscillatory signal decoding within the ERK cascade"

**SUPPLEMENTARY MATERIALS**

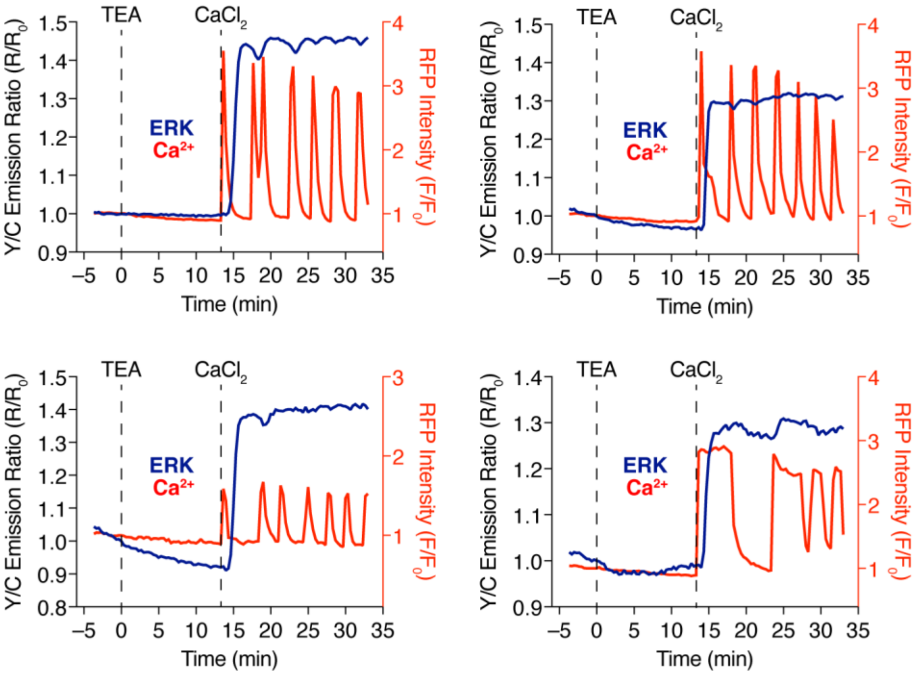


**Figure S1. Additional representative curves showing $Ca^{2+}$-dependent ERK activity in MIN6 cells – related to Fig. 1.** Single-cell timecourses from 4 additional TEA- and $CaCl_2$-treated MIN6 cells co-expressing EKAREV (ERK, dark blue curves) and RCaMP1d ($Ca^{2+}$, red curves) are shown.

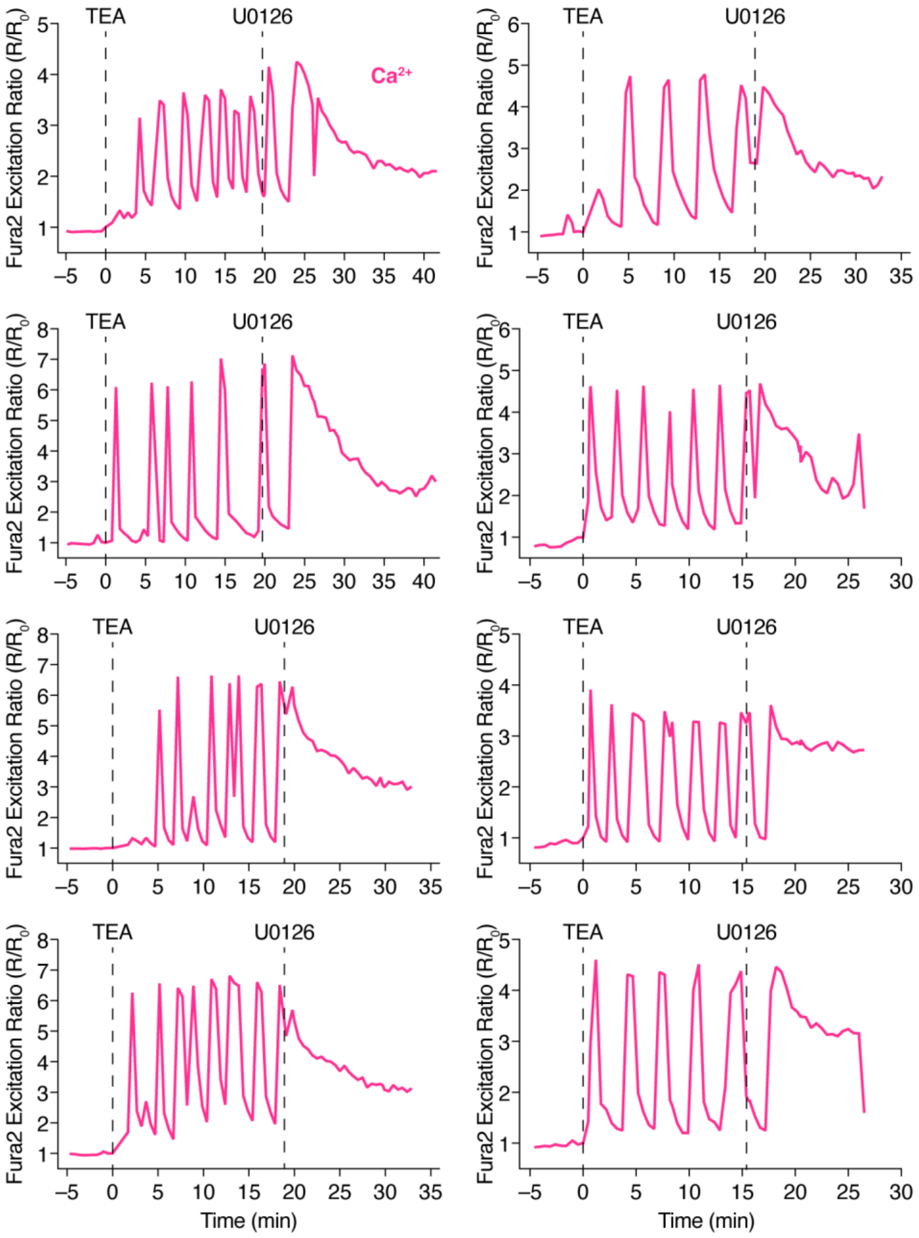


**Figure S2. Additional representative curves showing disruption of TEA-induced $Ca^{2+}$ oscillations via MEK inhibition with U0126 – related to Fig. 1.** Single-cell timecourses from 8 additional MIN6 cells stained with Fura-2 and sequentially treated with TEA and U0126 are shown.

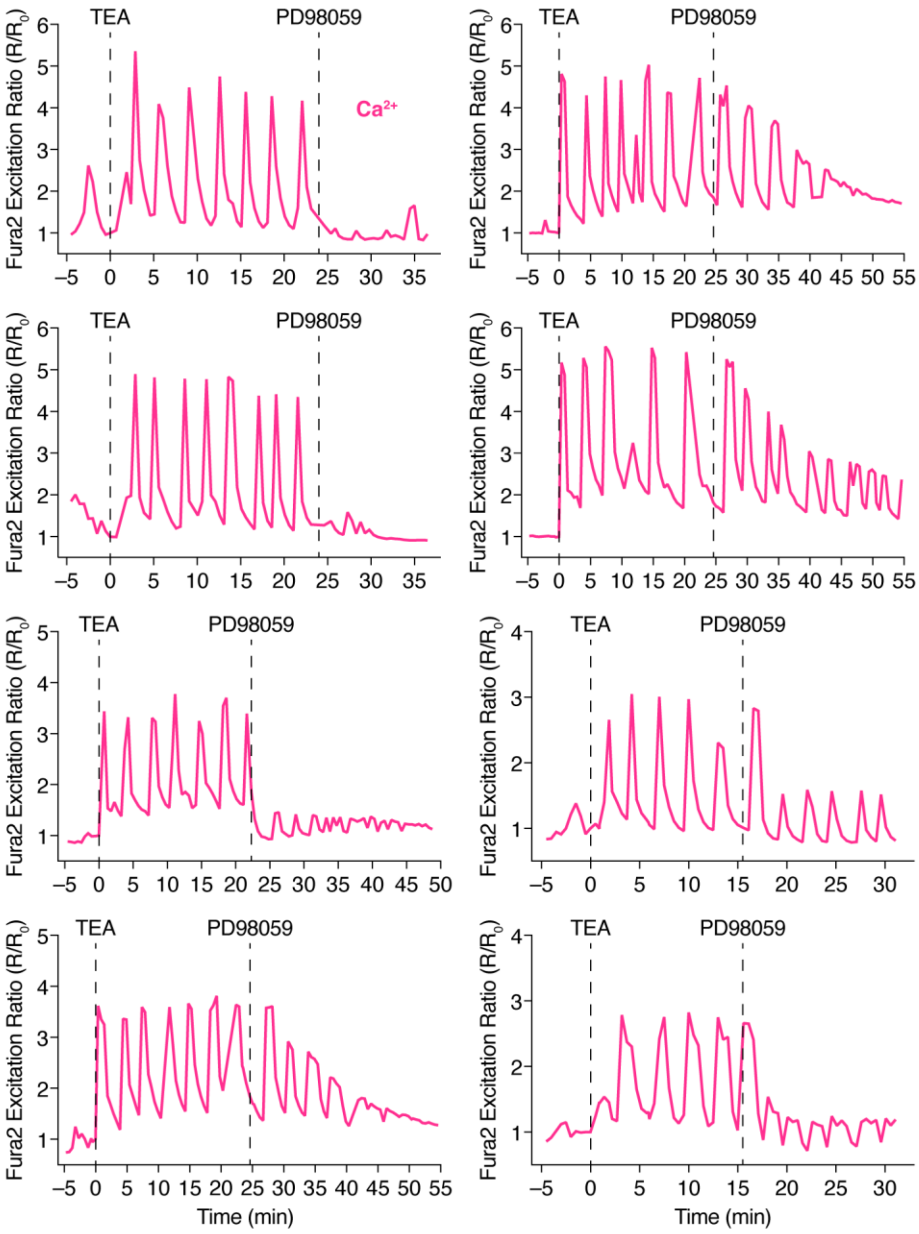


**Figure S3. Additional representative curves showing disruption of TEA-induced $Ca^{2+}$ oscillations via MEK inhibition with PD98059 – related to Fig. 1.** Single-cell timecourses from 8 additional MIN6 cells stained with Fura-2 and sequentially treated with TEA and PD98059 are shown.

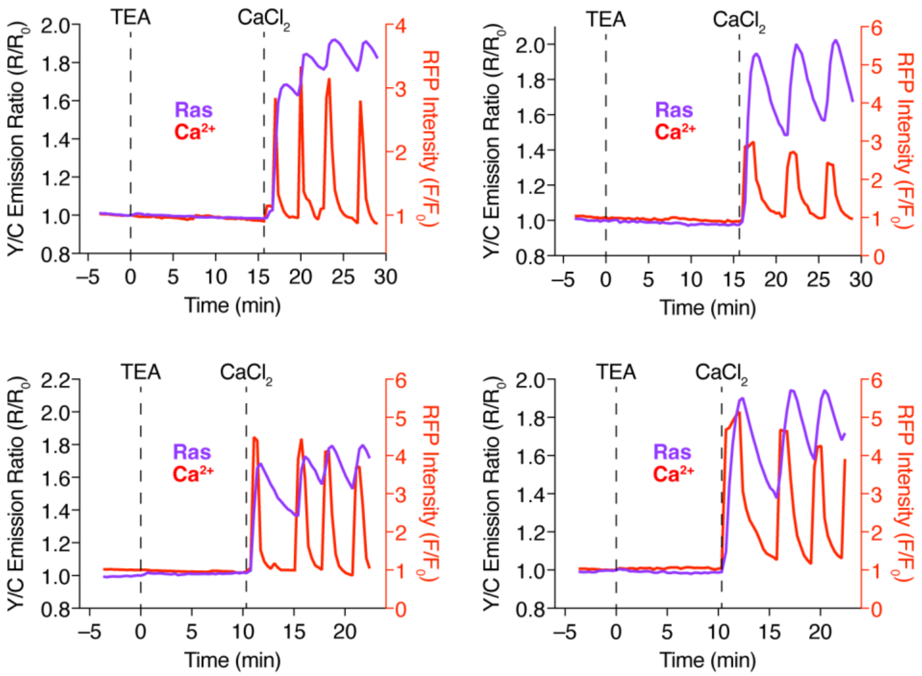


**Figure S4. Additional representative curves showing $Ca^{2+}$-dependent Ras activity in MIN6 cells – related to Fig. 2.** Single-cell timecourses from 4 additional TEA- and $CaCl_2$-treated MIN6 cells co-expressing RaichuEV-Ras (Ras, blue curves) and RCaMP1d ($Ca^{2+}$, red curves) are shown.

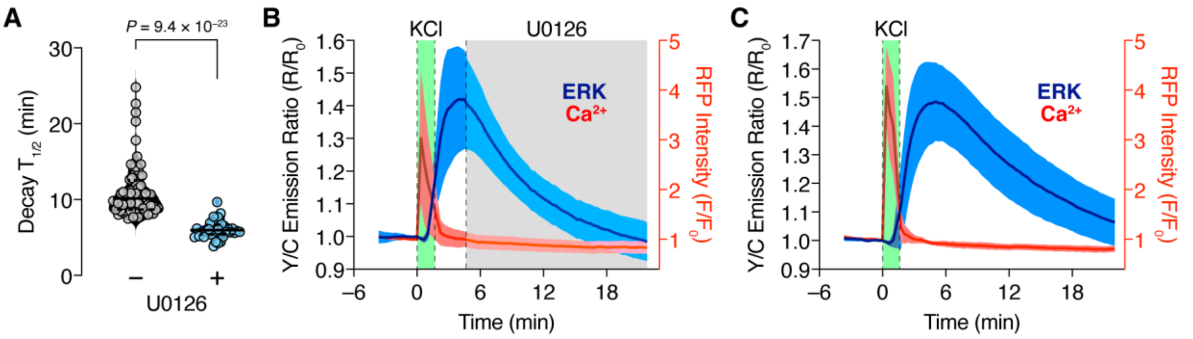


**Figure S5. MEK inhibition accelerates deactivation kinetics of $Ca^{2+}$-induced ERK response – related to Fig. 3.** (A) Summary quantification of the decay kinetics of EKAREV emission ratio response in MIN6 cells treated with a single, 1-min pulse or 30 mM KCl without (−, n = 70 cells from 4 independent experiments) or with (+, n = 36 cells from 3 independent experiments) 20 µM U0126 added at the peak of the EKAREV response. Data analyzed using Mann-Whitney U-test. Solid and dashed lines in violin plot indicate median and quartiles, respectively. (B, C) Representative average timecourses of the EKAREV emission ratio (ERK, dark blue curves) and RCaMP1d fluorescence intensity ($Ca^{2+}$, red curves) responses in KCl-stimulated MIN6 cells either (B) with or (C) without U0126 addition as indicated. Timecourses are representative of (B) 3 and (C) 4 independent experiments. Solid lines indicate the mean response; shaded areas, s.d.

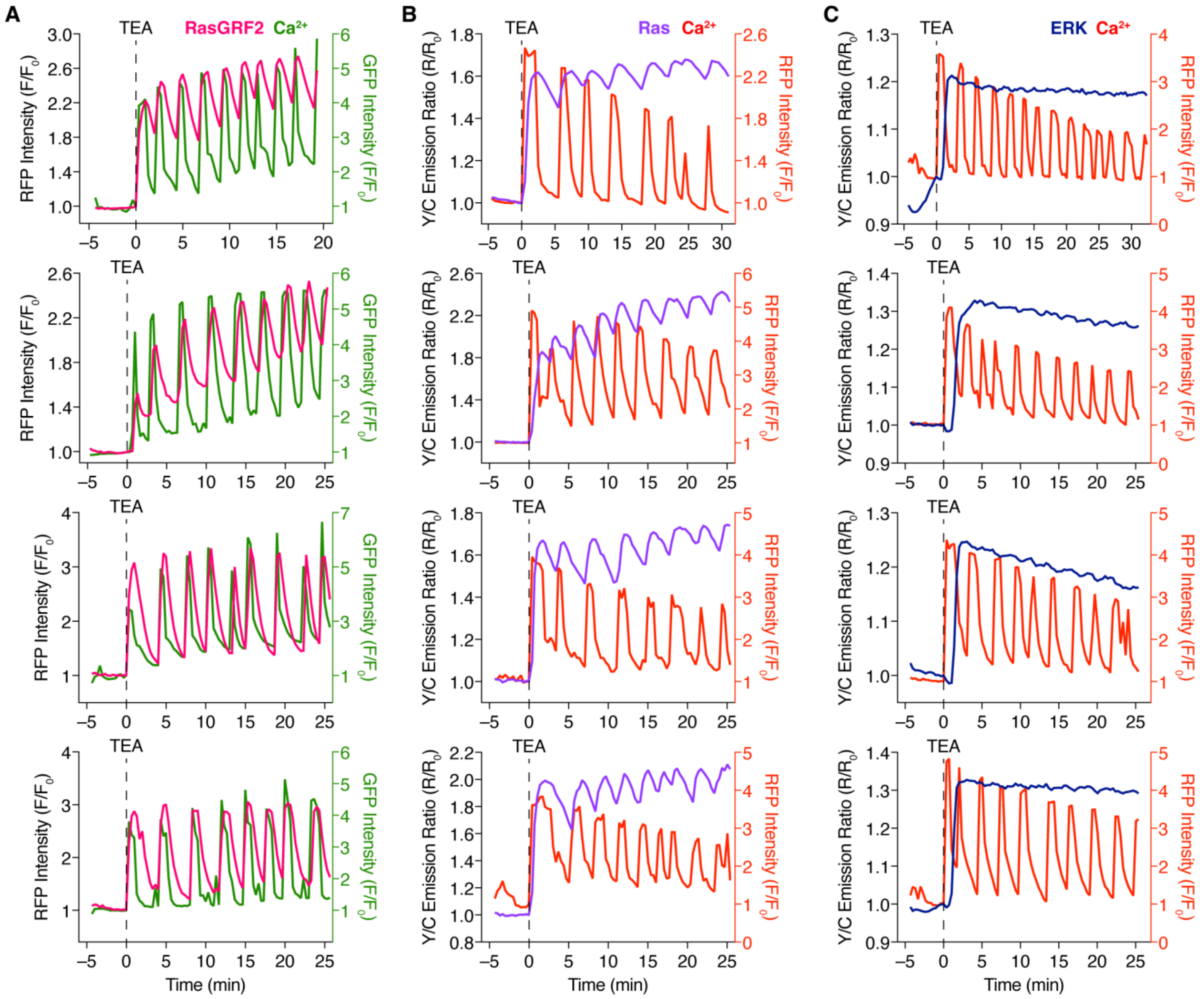


**Figure S6. Additional representative curves showing ERK cascade activity dynamics in response to $Ca^{2+}$ oscillations in MIN6 cells – related to Fig. 4.** Single-cell timecourses from TEA-stimulated MIN6 cells co-expressing (A) RasGRF2 (magenta curves) plus pmGCaMP3 (green curves), or either (B) RaichuEV-Ras (Ras, blue curves) or (C) EKAREV (ERK, dark blue curves) plus RCaMP1d ($Ca^{2+}$, red curves) from 4 additional cells each are shown.

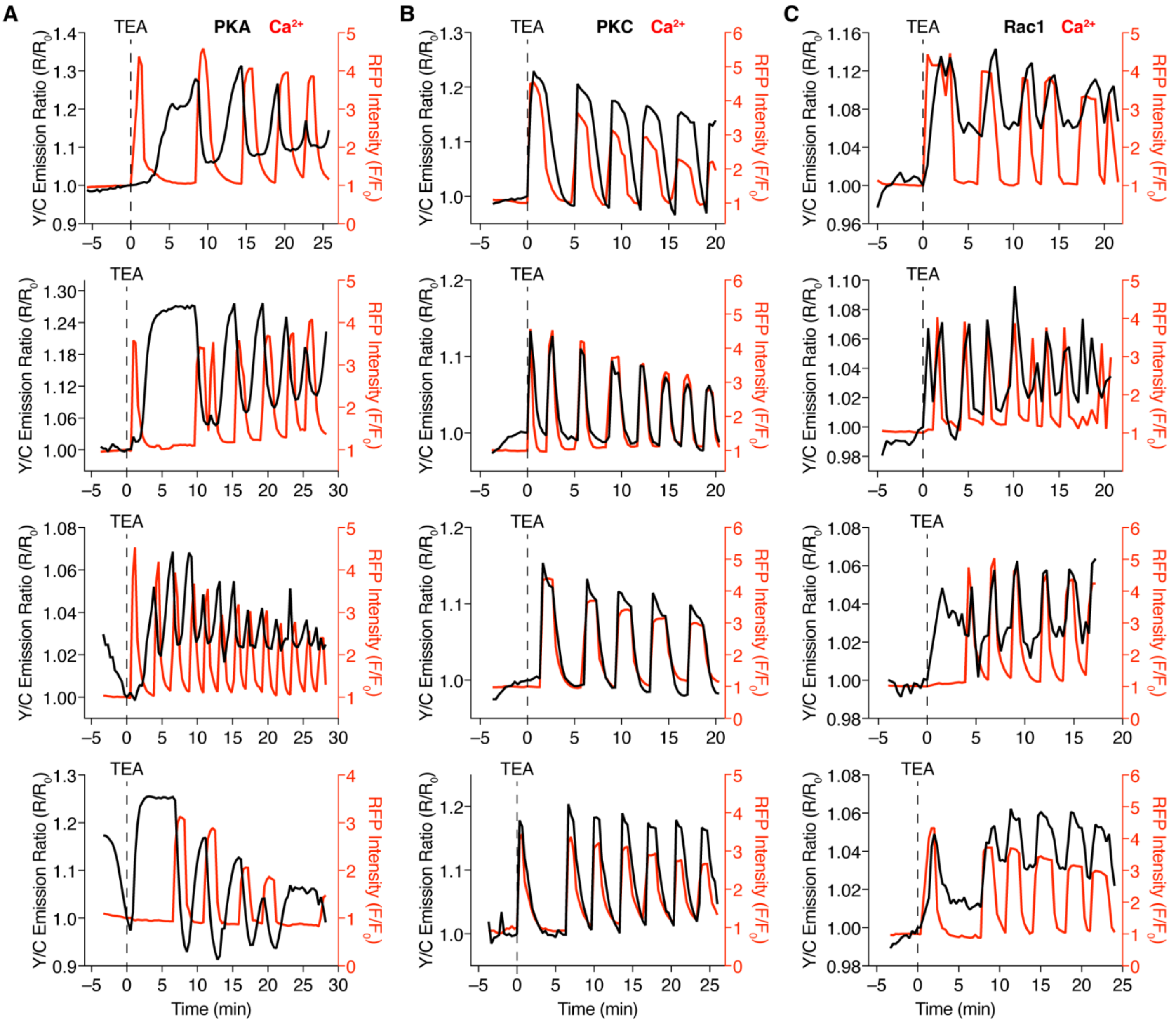


**Figure S7. Additional representative curves showing PKA, PKC, and Rac1 activity dynamics in response to $Ca^{2+}$ oscillations in MIN6 cells – related to Fig. 5.** Single-cell timecourses from TEA-stimulated MIN6 cells co-expressing (A) AKAR4, (B) CKAR2, or (C) RaichuEV-Rac1 (black curves) plus RCaMP1d (red curves) from 4 additional cells each are shown.
